# Frequency of a nucleotide overhang at the 5’ end of hemorrhagic fever mammarenavirus genomes in public sequence data

**DOI:** 10.1101/2022.02.04.479072

**Authors:** Yutaro Neriya, Shohei Kojima, Mai Kishimoto, Arata Sakiyama, Takao Iketani, Tadashi Watanabe, Yuichi Abe, Hiroshi Shimoda, Keisuke Nakagawa, Takaaki Koma, Yusuke Matsumoto

## Abstract

Mammarenaviruses, such as Lassa virus and South American hemorrhagic fever (SAHF) virus, cause severe hemorrhagic fevers in humans, and pose major threats to public health. Mammarenaviruses consist of a bi-segmented negative-sense RNA genome in which the 5’ and 3’ ends form complementary strands that serve as a replication promoter. Some mammarenaviruses have a nucleotide overhang at the 5’ genome end. By examining the complementarity of 5’ and 3’ genome ends using public mammarenavirus genome sequences, we found that the 5’ guanine overhang (5’-G overhang) was present more frequently in Lassa and SAHF viruses than in other viruses. The 5’-G overhang in the Lassa and SAHF virus sequences was found to be restricted to the L and S segments, respectively. If the genome end sequence data in the public database are accurate, the 5’-G overhang may be related to the high pathogenicity of mammarenaviruses in humans.

## Introduction

Arenaviruses (genus *Mammarenavirus*, family *Arenaviridae*, order *Bunyavirales*) are divided into Old World and New World complexes (1). The most significant pathogen in the Old World complex is Lassa virus, the cause of hemorrhagic Lassa fever in humans. Lymphocytic choriomeningitis virus (LCMV) is also an Old World arenavirus that can cause a serious neurological disease called lymphocytic choriomeningitis, especially in prenatal and immunocompromised humans (2). The New World arenaviruses are divided into three major clades (A, B, and C), with clade B containing the Junín, Machupo, Guanarito, and Sabiá viruses, which cause Argentinian, Bolivian, Venezuelan, and Brazililian hemorrhagic fevers, respectively; collectively, these are referred to as South America Hemorrhagic fever (SAHF) (3). The mammarenavirus genome consists of two negative-strand RNA segments encoding four viral proteins: a large (L) RNA-dependent RNA polymerase (RdRp) and a matrix protein (Z) within the large RNA segment (L segment), and a nucleocapsid protein (NP) and a glycoprotein precursor (GPC) within the small RNA segment (S segment). The genomic RNA is encapsidated by NP and bound at both genome ends by L protein to form pseudocircularized ribonucleoprotein complexes (RNPs) (4–6). The genomic RNPs are enveloped by a lipid bilayer decorated with mature viral glycoproteins to form infectious virions.

In mammarenaviruses, the genome ends exhibit terminal complementarity of the 5’ and 3’ ends in each of the L and S segments to form a replication promoter. The interaction of each 5’ and 3’ end to L protein is essential for the initiation of RNA synthesis (7,8). The 5’ end of some mammarenavirus genomes contains an additional nontemplated guanine residue (5’-G overhang) (9–12), which has been considered to be generated by a prime-and-realign mechanism. During mammarenavirus replication, it is thought that RdRp initiates at position +2 of the promoter to generate a GC dinucleotide primer that realigns, leading to base pairing between the C and a G at position +1 of the template (prime-and-realign) (13). Subsequent elongation by RdRp results in the synthesis of antigenomic RNPs with a pseudotemplated G residue at the 5’ end (5’-G overhang). The antigenomic RNPs are used as templates for mRNA synthesis, and the GPC and Z mRNAs are transcribed from the S and L segments, respectively. The antigenomic RNPs also serve as templates for GC-primed replication to produce progeny genomic RNPs with 5’-G overhang. The mammarenavirus replication has been also shown to be initiated with CG primer that starts with precise genomic end (+1 position) without generating 5’-G overhang (13). It remains unclear whether the initiation of RNA synthesis and the genomic terminal structures are common between all mammarenaviruses, and the biological significance of the 5’-G overhang has not been studied in detail.

In this study, we constructed a list of mammarenavirus promotors by using public sequence data to characterize the promoter structures of various species of mammarenaviruses. We examined the conservation of sequences of promoters among mammarenaviruses, and analyzed the genomic sequences for the presence of 5’- and 3’-overhangs.

## Methods

### List of mammarenavirus promoters

Forty-six mammarenavirus species were registered on the International Committee on Taxonomy of Viruses (ICTV) list (https://talk.ictvonline.org/) on December 7th, 2021. The complete sequences of S and L segments of mammarenaviruses available on the National Center for Biotechnology Information (NCBI) associated with Genbank accession numbers as listed in Supplementary Table 1 were used for the analysis. The sequences of the extreme 40 nucleotides (nts) of each of the 5’ and 3’ ends of mammarenaviruses were collected, and the complementarity between the 5’ and 3’ ends of the sequences was examined. The counts of G:C and A:U complementarity and each of the nts (A, U, G, and C) in the promoter region are shown in Supplementary Table 1. The conservation of the nts in the promoter was analyzed by using a sequence logo generator, Weblogo (https://weblogo.berkeley.edu/logo.cgi).

### Analysis of the 5’- overhang in the sequences of the *Arenaviridae* family

Sequences annotated as *Arenaviridae* were downloaded from the NCBI refseq database on January 7th, 2022. There were 5,274 *Arenaviridae* sequences in the database. Sequences shorter than 1,000 nt (n = 2,343) were omitted from this analysis. We first checked for the presence of the promoter sequence, CGCA, in the 8-nt ends of the sequences. The promoter sequence was present in both ends of 502 sequences. Then, the overhangs were detected from the 502 sequences by a custom Python script. The codes used for this analysis is available on GitHub (https://github.com/shohei-kojima/Arenaviridae_overhang_analysis_2022).

## Results and Discussion

### Construction of a list of promoters in mammarenaviruses

To examine the conservation of the promoter sequence among virus species, we compared the both extreme 40 nts of the mammarenavirus genome sequences registered on the ICTV list. Some mammarenavirus sequences available on the NCBI database did not have complete genomic terminal information (14,15). Sequences that showed complementarity between +1 and +4 within 0 to 2 nt shifting of the 5’ or 3’ ends for both L and S segments were selected (Figure 1A). We obtained 23 mammarenavirus promoter sequences of both the L and S segments, as listed in Supplementary Table 1 (shown as the positive-strand antigenomic forms). The conservation of the extreme 38 nts (without the overhang) of each of the 5’ and 3’ ends of the L and S segments among the 23 virus species was analyzed by using a sequence logo generator, Weblogo (Figure 1B). The first 5 nts of the 5’ and 3’ ends were completely conserved in both the L and S segments in the 23 mammarenaviruses. Although the +6 nt in the 3’ end varied in the L segment, that in the S segment was U in almost all samples. This difference may result in differences in the promoter strength of the L and S segments to create these segments in a different ratio in infected cells as proposed in other segmented RNA viruses (16,17). To further characterize the promoter structure, the counts of G:C and A:U complementarity and A, U, G, and C in the first 40 nts (for 38 to 39 nts in the opposite strand of 2-nt and 1-nt overhangs, respectively) of the L and S segments were determined (Figure 1C and Supplementary Table 1). G:C complementarity was significantly higher than A:U complementarity in both the L and S segments; this result differed from that of tri-segmented viruses belonging to the order *Bunyavirales*, which showed higher A:U complementarity in the promoters (18). The L and S segments contained significantly less A than U, G, and C in the 5’ ends, which results in an nt bias between the genomic and antigenomic promoters. This may function to regulate the replication balance between genome and antigenome to produce different ratios of various viral RNA species as observed in infected cells (19). The 5’-CGCA-UGCG-3’ sequence was the most conserved genome end sequence in the 23 mammarenaviruses (Figure 1D, non-overhang). The 5’-**<u>G</u>**CGCA-UGCG-3’ sequence showed the presence of an unpaired 5’ guanine nt (Figure 1D, 5’-G overhang). Among the 23 viruses, the 5’-G overhang was found in the Lassa, Guanarito, Junín, Machupo, Sabiá, and Oliveros viruses (Figure 1D), and the 2-nt 3’-overhang was found only in the Lunk virus S segment (Supplementary Table 1) in the public database. All viruses, except for Oliveros virus (20,21), with the 5’-G overhang are associated with human hemorrhagic fever, and are categorized as biosafety level 4 pathogens.

**Figure 1.**
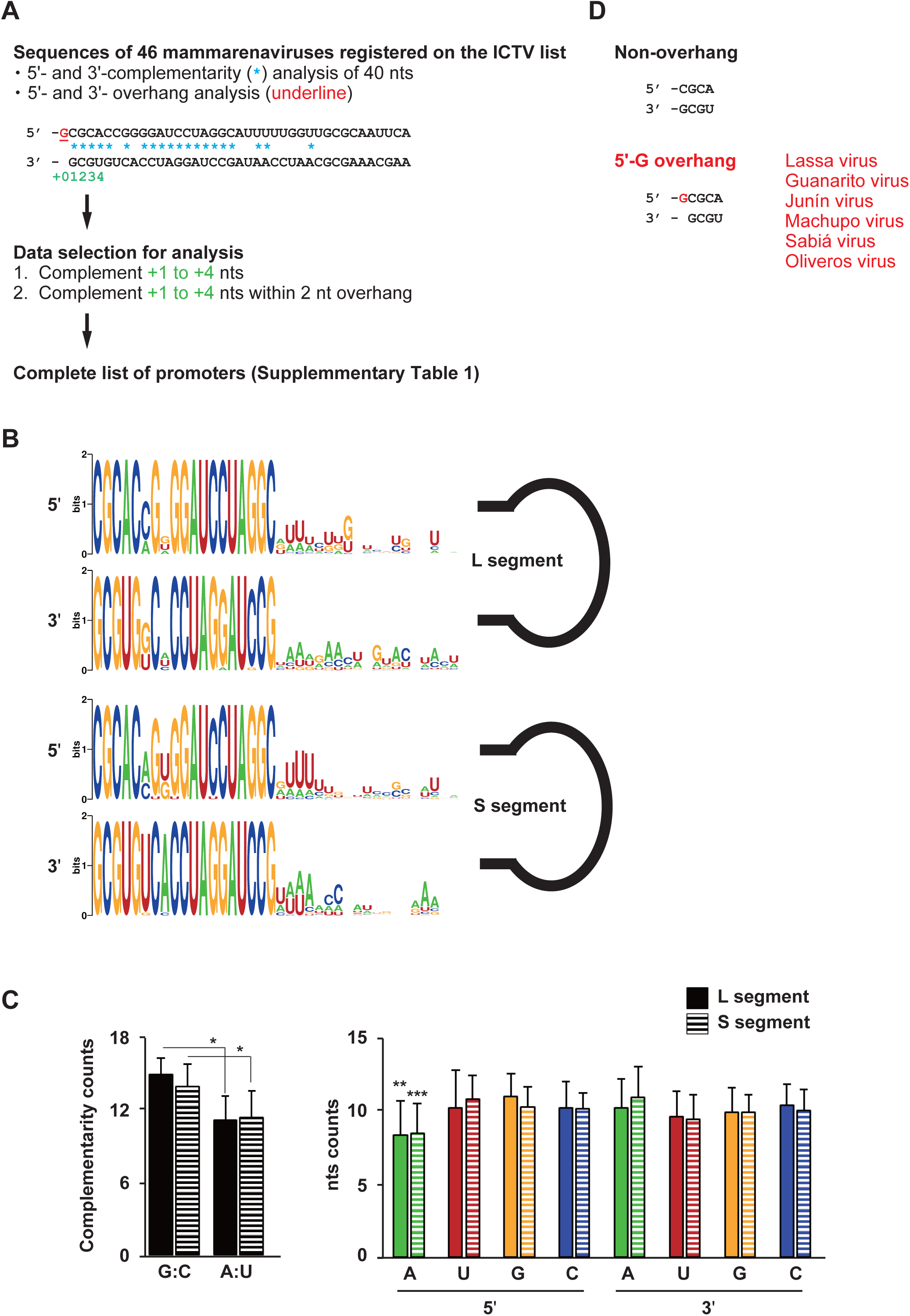
Comprehensive analysis of the genomic promoters in mammarenaviruses. (A) Schematic diagram of mammarenavirus promoter sequences and the complementarity of the 5’ and 3’ ends. (B) The sequence conservation of the 38 nts in the 5’ and 3’ genome ends of the L and S segments (non-overhang region) among 23 mammarenavirus species was analyzed by a sequence logo generator, Weblogo. (C) Counts of G:C and A:U complementarity and each nt in the first 40 nts. Bars represent the means and statistical deviations. Inferential statistical analysis was performed by one-way analysis of variance (ANOVA) followed by the Tukey’s test. *p < 0.01, **A > U, G, and C in 5’ end (p < 0.05), *** A > U, G, and C in 3’ end (p < 0.05). (D) Schematics of the first 4 to 5 nts in the non-overhang and 5’-G overhang genome ends. Virus species possessing the 5’-G overhang are shown. All sequences are shown as the positive-strand antigenomic form.

### Analysis for the 5’-G overhang in the *Arenaviridae* family using public sequence data

To expand the study of viruses having the 5’-G overhang, we next analyzed sequences with 5’-CGCA-UGCG-3’ (non-overhang), 5’-**<u>G</u>**CGCA-UGCG-3’ (5’-G overhang), or 5’-**<u>(N)N</u>**CGCA-UGCG**<u>N(N)</u>**-3’ (1- or 2-nt overhang [not the 5’-G overhang]) in sequences of the *Arenaviridae* family (Figure 2A). Among the virus sequences, artificially generated sequences (e.g., generated by reverse genetics virus engineering) were excluded. We found that 345 sequences did not have the overhang (5’-CGCA-UGCG-3’), whereas 58 sequences had the 5’-G overhang (5’-**<u>G</u>**CGCA-UGCG-3’; Supplementary Table 2). The sample numbers of over 5 sequences per virus species are shown in Figure 2B. We found that virus genomes with the 5’-G overhang were restricted to the Lassa, LCMV, Junín, Machupo, Guanarito, Sabiá, and Oliveros viruses (Figure 2B and C). All LCMVs with the 5’-G overhang found in this study were clones obtained from virus-inoculated laboratory mice (22,23). We next determine which segments (L or S) contained the 5’-G overhang in the Lassa, Junín, Machupo, Guanarito, Sabiá, and Oliveros viruses. Guanarito virus sequences had the 5’-G overhang only in the L segment, Lassa and Oliveros virus sequences had the 5’-G overhang only in the S segment, and the other sequences had it in both segments (Figure 2D and E). L segments with the 5’-G overhang were found only in the New World viruses causing SAHF. In general, the Old World viruses are geographically confined to the African continent, with the exception of LCMV that circulates globally. The New World viruses are found on the South and North American continents. Regardless of the region where the viruses are distributed, the 5’-G overhang was found specifically in hemorrhagic fever virus sequences on both continents. Although Oliveros virus is a New World virus, it had the 5’-G overhang in the L segment, unlike other SAHF viruses. This slight difference in the Oliveros virus from other SAHF viruses might be the reason for its non-pathogenic characteristic.

**Figure 2.**
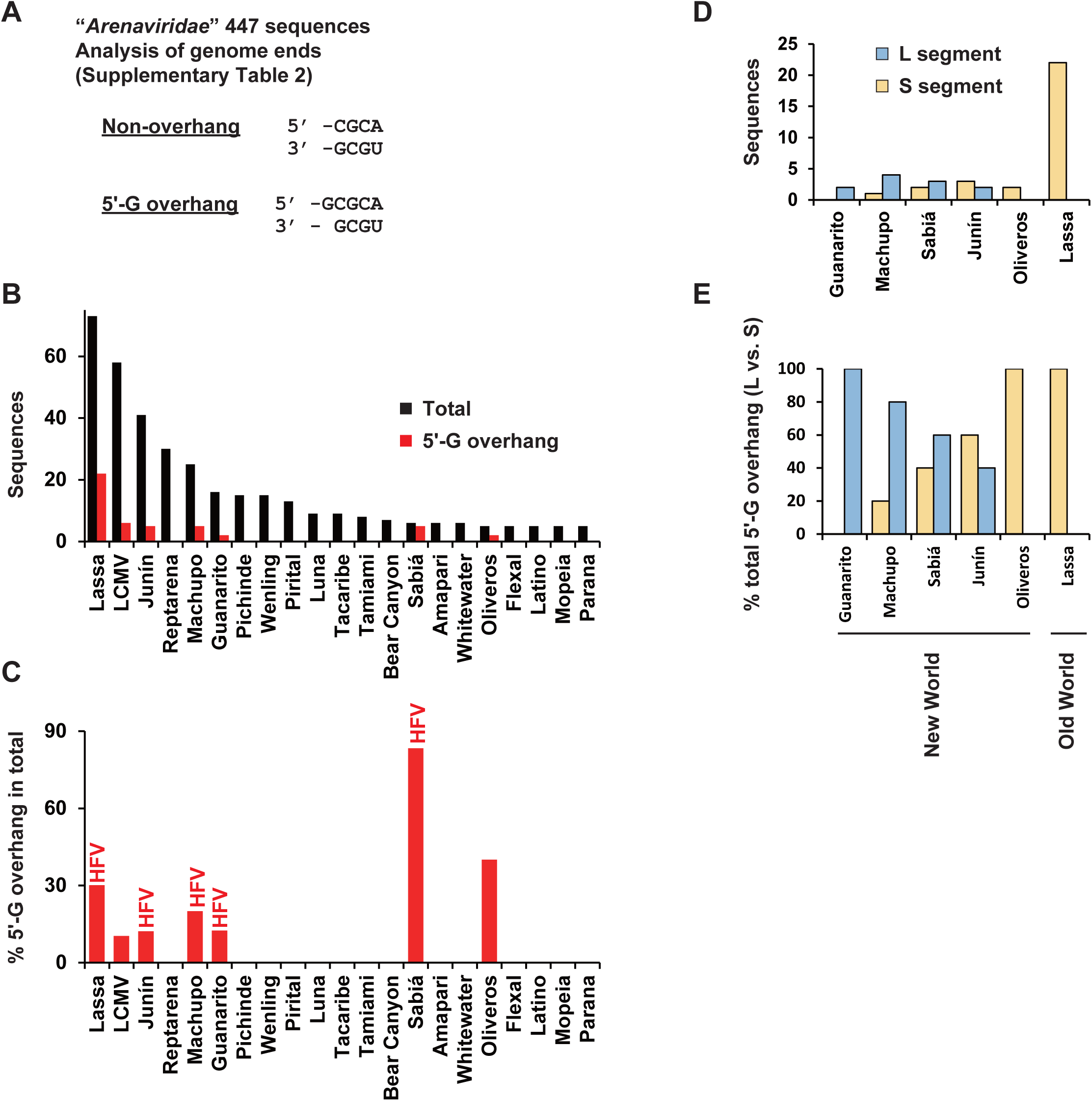
Analysis of the overhang at the genome end in the *Arenaviridae* family. (A) Schematic diagram of the analysis of genome end sequences in the *Arenaviridae* family. Viral antigenome sequences starting with 5’-CGCA and ending with UGCG-3’ were recognized as not having the overhang (non-overhang), and those starting with 5’-GCGCA and ending with UGCG-3’ were recognized as having the 5’-G overhang. (B) The number of sequences per virus species recognized as not having (non-overhang) or having the 5’-G overhang shown in panel A. (C) The percentage of sequences possessing the 5’-G overhang among the total sequences. HFV: hemorrhagic fever virus. The full virus names are shown in Supplementary Table 2. Reptarena: unidentified reptraena virus strains. (D) The number of sequences possessing the 5’-G verhang in the L and S segments. (E) The percentage of sequences possessing the 5’-G overhang in the L and S segments (L + S = 100%).

### Minor genome end sequences in the database

Among the 447 sequences described in Figure 2A, we found 35 sequences that did not have 5’-CGCA-UGCG-3’ (non-overhang) or 5’-**<u>G</u>**CGCA-UGCG-3’ (5’-G overhang). Two minor non-overhang sequences, i.e., 5’-**<u>G</u>**CGCA-UGCG**<u>C</u>**-3’ and 5’-**<u>U</u>**CGCA-UGCG**<u>A</u>**-3’, were found in some viruses, including biosafety level 4 pathogens (5’-**<u>G</u>**CGCA-UGCG**<u>C</u>**-3’ in Lassa, Lujo, and Junín viruses, and 5’-**<u>U</u>**CGCA-UGCG**<u>A</u>**-3’ in Chapare virus; Figure 3A). These sequences may have evolved from the 5’ overhang with an additional nt on the opposite end of the overhang. We also found some variations in the 5’ overhang (not the 5’-G overhang; Figure 3B).The 5’-**<u>CG</u>**CGCA-UGCG**<u>C</u>**-3’ sequence found in the Lassa and Junín viruses seemed to be a 1-nt addition to the 5’ end of the minor non-overhang shown in Figure 3A. The 3’-C overhang (5’-CGCA-UGCG**<u>C</u>**-3’) was found in the Lassa, Oliveros, and other virus sequences (Figure 3B). When viral replication occurs using the genome with a 5’-G overhang as a template, the 3’ end of the newly replicated RNA terminates with a C complementary to the overhanging G. It has been presumed that mammarenaviruses remove this last C to retain genome integrity (13). The 3’-C overhang may represent this C that was not removed during the replication process. The 3’ overhang (5’-**<u>G</u>**CGCA-UGCG**<u>CG-</u>**3’) found in Lassa virus (Figure 3C) was also considered to have arisen from a minor non-overhang. There were a few virus sequences possessing a 2-nt overhang both in the 5’ and 3’ ends, i.e., an unidentified *arenavirus* sp. and Lunk virus (Figure 3B and C). The mechanism of their generation and their biological properties remain unknown.

**Figure 3.**
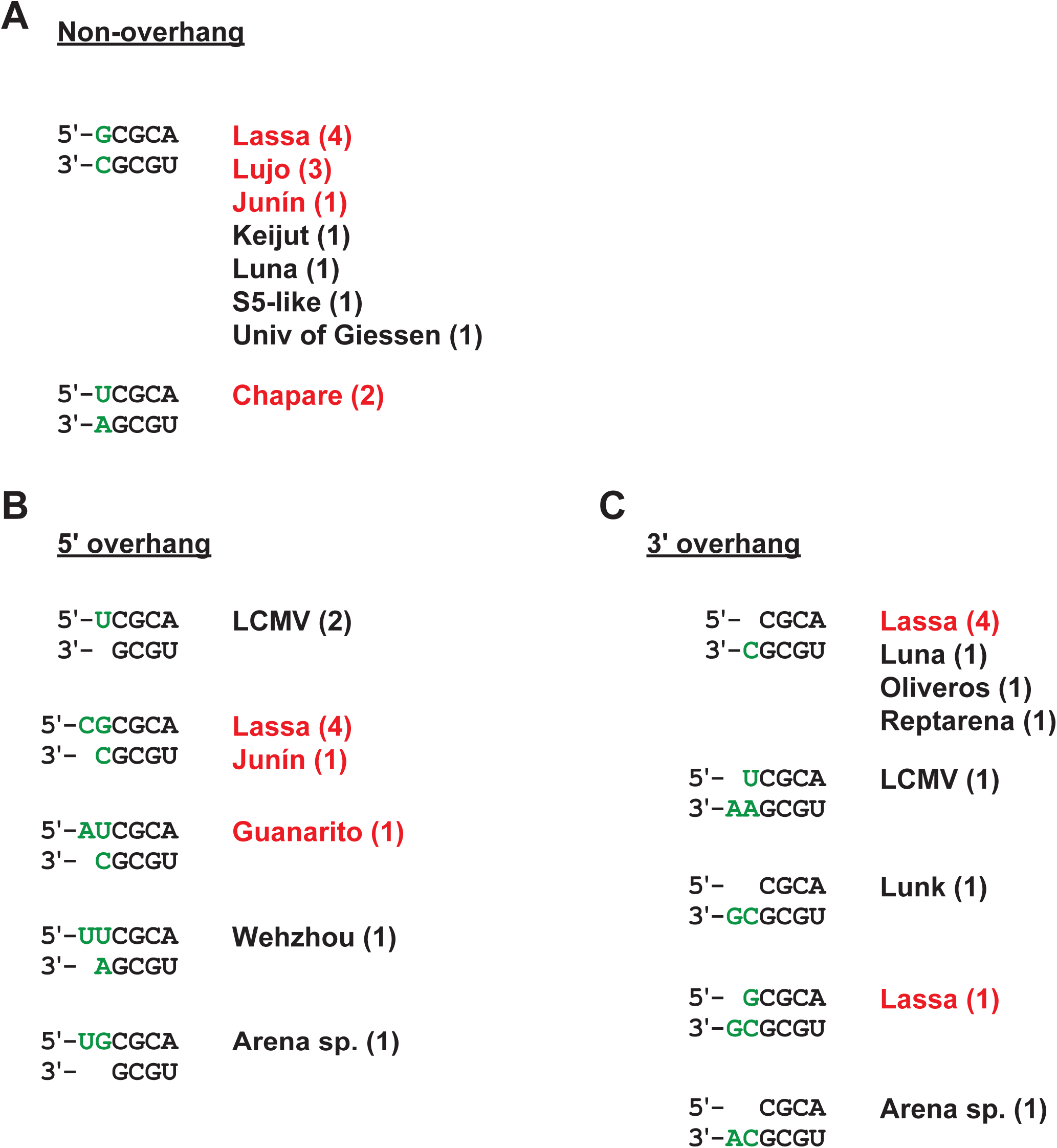
Minor genome end sequences in the *Arenaviridae* family. (A) Minor sequences without the overhang (non-overhang). (B) Minor sequences with the 5’ overhang. (C) Sequences with the 3’ overhang. Additional nts outside of the 5’-CGCA-UGCG-3’ sequence are shown in green. HFVs are shown in red.

### Dependence on the accuracy of the sequence database

All of the results of this study depend on the accuracy of the public genome sequences. If the genome end sequences in the NCBI database were correct and accurately registered, hemorrhagic fever viruses appear to have a higher frequency than other non-pathogenic arenaviruses of containing the 5’-G overhang in their genomes. However, some of the sequences in the database were obtained from full-genome amplification via reverse transcription polymerase chain reaction, and in those cases, the extreme sequences are derived from oligonucleotide primers. Further experiments are needed to validate whether the 5’-G overhang correlates with pathology. Even if a mutant arenavirus that has an artificially attached or removed 5’-G overhang is produced using reverse genetics, it would be difficult to evaluate whether changes in its pathogenicity are due to the presence or absence of the 5’-G overhang, because the virus can automatically synthesize the overhang by the prime-and-realign mechanism of the L protein. Identification of the functional region in the L protein that is involved in the prime-and-realign mechanism and that confers a 5’-G overhang is essential to prove a correlation between the 5’-G overhang and viral pathogenicity.

### The advantage of the 5’-G overhang for arenavirus growth and pathogenesis

The high frequency of the 5’-G overhang in hemorrhagic fever virus sequences in the public database suggests that hemorrhagic fever viruses have a higher ability than other viruses to produce the overhang via prime-and-realign. It was hypothesized that the 5’-G overhang might increase the pathogenicity of these viruses in humans. It has been proposed that the 5’-G overhang in the promoter protects this portion of double-stranded RNA from being recognized by retinoic acid-inducible gene I (24). The presence of the 5’-G overhang was shown to be important for stimulating the RNA synthesis of purified Machupo and Lassa virus L protein (25,26). These facts indicate that the presence of the 5’-G overhang may be an important factor associated with viral pathogenicity via the enhancement of host immune evasion and polymerase activity.

## Author contributions

Y.N., S.K., M.K., A.S., T.I., T.W., Y.A., H.S., K.N., T.K. and Y.M. conceived and designed the study, performed the analyses, analyzed the data. Y.M. wrote the manuscript. All authors have read and agreed to the manuscript.

